# Tiled STED Imaging of Extended Sample Regions

**DOI:** 10.1101/789487

**Authors:** Jonatan Alvelid, Ilaria Testa

## Abstract

Stimulated Emission Depletion (STED) nanoscopy has become one of the most used nanoscopy techniques over the last decade. However, most recordings are done in specimen regions no larger than 10–30 × 10–30 µm^2^ due to aberrations, instability and manual mechanical stages. Here, we demonstrate automated STED nanoscopy of extended sample regions up to 0.5 × 0.5 mm^2^ by using a back-aperture-stationary beam scanning system. The setup allows up to 80–100 x 80–100 µm^2^ field of view (FOV) with uniform spatial resolution, a mechanical stage allowing sequential tiling to record larger sample areas, and a feedback system keeping the sample in focus at all times. Taken together, this allows automated recording of theoretically unlimited-sized sample areas and volumes, without compromising the achievable spatial resolution and image quality.

## 1. Introduction

STED nanoscopy[1] is widely used in biological applications[2] thanks to its flexibility in accessing fixed and live samples with multi-colour[3] and optical sectioning abilities. The technique belongs to the family of coordinate-targeted nanoscopy methods[4] together with RESOLFT[5–7], and overcomes the diffraction limit by using the inherent stimulated emission process of most fluorescent molecules to keep them non-detectable in space and time. This is typically implemented by using a Gaussian-shaped excitation beam and a red-shifted, phase-shaped depletion beam. By imparting a helical phase shift on the beam in a conjugate back focal plane of the objective lens, the beam forms a doughnut-shaped point spread function (PSF) at the focal plane, containing a central zero. The phase shift is either produced using a phase retarding plate, as in the earliest STED implementation[8], or more recently with a spatial light modulator (SLM)[9]. The SLM has the additional benefit of being electronically controlled, allowing simpler alignment procedures[10], 3D STED pattern generation[11] as well as aberration correction[12].

In order to obtain the best possible STED image, two parameters are of utmost importance and must be optimal across the whole FOV. Firstly, the phase profile of the depletion beam, translating to the quality of the zero and the homogeneity of the doughnut to achieve isotropic depletion in the focal plane. Secondly, the co-alignment of the excitation maxima and the minima of the depletion beam, no matter if the depletion beam is a doughnut for 2D STED, top hat for 3D STED, or another shape. Both are responsible for the spatial resolution and the fluorescence signal in the final STED image. Optical aberrations can therefore influence the STED image quality by compromising the PSF of the excitation and STED beam. This becomes increasingly important when beam-scanning large FOVs where off-axis aberrations and vignetting can occur.

Additionally, chromatic aberrations often affect the performance of a STED microscope, as the excitation and depletion beams are inherently of different wavelengths. To minimize or correct chromatic aberrations, multiple approaches have been developed such as easySLM-STED[13], which uses an SLM to re-align the excitation and STED beam at the border of the FOV, achieving a maximum tile size of 20 × 20 µm^2^ combined into a final FOV of 100 × 100 µm^2^. Another way to possibly overcome the chromatic aberrations is by preventing any chromatism in the system, such as in two-photon excitation STED with a single wavelength[14]. Here, we show that it is possible to achieve an extended FOV with minimal aberrations by using optical elements and a scanning system that generates stationary beams in the back aperture by using an achromatic spherical mirror instead of a lens. This results in a system with co-aligned beams all the way out to at least 40 µm from the centre, and ultimately the possibility to record FOVs as large as 80 × 80 µm^2^ without any significant degradation in resolution at the outermost positions. An alternative scanning approach with similar performance is the quadscanner[15].

In addition to extending the FOV to 80–100 × 80–100 µm^2^, our system also features tile-scanning to access even larger specimen regions. This is done by stitching together multiple images recorded sequentially in time and space with a motorized mechanical scanning system in a mosaic fashion. The tile scan is stabilized thanks to a feedback loop-based focus lock system ensuring that the focal plane is always kept the same during recordings. Importantly, tile-scanning STED is automated with a custom-developed and open-source Python software, which makes the system robust and easy to use.

Finally, our tile-scanning STED can record theoretically unlimited-sized extended sample regions, thus allowing the technique to be leveraged in a large range of biological studies ranging from sub-cellular analysis to entire cells and even organs.

## 2. Material and Methods

### 2.1 STED set-up

All images were acquired on a custom-built STED setup. Excitation of the dyes was done with two pulsed diode lasers, one at 561 nm (PDL561, Abberior Instruments, Göttingen, Germany), and one at 640 nm (LDH-D-C-640, PicoQuant, Berlin, Germany), both with pulse widths of ∼60 ps. The 561 nm and 640 nm excitation laser beams are coupled into a fibre (P5-488PM-FC-2, Thorlabs, Newton, NJ, USA) in order to co-align the beams and shape the wave fronts. The 775 nm depletion laser beam (KATANA 08 HP, OneFive GmbH, Regensdorf, Switzerland), with a pulse width of 530 ps, is led through an AOM (MT110-B50A1.5-IR-Hk + MDS1C-B65-34-85.135-RS, AA Opto Electronic, Orsay, France) for line-by-line power modulation and then coupled into a polarization-maintaining single-mode fiber (PMJ-3AHPM3S-633-4/125-3-3-1, OZ Optics, Ottawa, ON, Canada). The two orthogonal polarizations are split up in an interferometer, slightly time shifted and then co-aligned again. The two orthogonally polarized beams are then subsequently shaped using a vortex and a top-hat phase mask respectively on a spatial light modulator (LCOS-SLM X10468-02, Hamamatsu Photonics K.K., Hamamatsu, Japan), which can spatially phase shift the light between 0–2π on a pixel-by-pixel basis. The SLM only affects a single polarization direction, hence we can separately shape the orthogonally polarized beams by turning the polarization in between using a double pass of a λ/4 wave plate. The excitation and depletion beams are merged into a single beam path through two dichroic mirrors, passing through a λ/4 wave plate and a λ/2 wave plate to create the circular polarization necessary for optimal depletion focus formation and subsequently scanned using fast galvanometer mirrors for the fast and slow scanning axis (galvanometer mirrors 6215H + servo driver 71215HHJ 671, Cambridge Technology, Bedford, MA, USA). Using a concave mirror between the galvanometer mirrors and the scan and tube lens, the respective galvanometer mirror planes are both conjugate to the pupil plane of the objective, ensuring minimal distortion aberrations in the sample plane. The objective used is a 100x/1.4 oil immersion objective (HC PL APO 100x/1.40 Oil STED White, 15506378, Leica Microsystems, Wetzlar, Germany). The fluorescence is collected back through the same beam path, and is de-coupled from the incoming laser beams through a multi-bandpass dichroic mirror (ZT405/488/561/640/775rpc, Chroma Technology, Bellows Falls, VT, USA) and a long-pass dichroic mirror (ZT514RDC, Chroma) after being de-scanned by the galvanometer mirrors. The combined fluorescence signal is then passed through a pinhole of size 75 µm (1.28 Airy disk units) and separated with another dichroic mirror (Di02-R635, Semrock, Rochester, NY, USA) into two channels. Channel 1 (red) has a notch filter (NF03-785E-25, Semrock), a bandpass filter (ET705/100m, Chroma), and the fluorescence is focused onto a free space APD (SPCM-AQRH-13-TR, Excelitas Technologies, Waltham, MA, USA). Channel 2 (green) has a notch filter (ZET785NF, Chroma), a bandpass filter (ET615/30m, Chroma), and the fluorescence is focused onto a 62.5 µm core diameter multi-mode fibre (M31L01, Thorlabs) coupled to an APD (SPCM-AQRH-14-FC, PerkinElmer, Waltham, MA, USA). The fluorescence signal is collected by a NI-DAQ acquisition board (PCIe-6353, National Instruments, Austin, TX, USA), also used to control the lasers and the scanning.

**List of components: Lenses**: L1: 300 mm, L2: 200 mm, L3: 100 mm, L4: 200 mm, L5: 400 mm, L11: 150 mm, L13: 200 mm, L14: 200 mm (all AC254-XXX-B-ML, Thorlabs), L6: 100 mm, L7: 200 mm, L8: 200 mm, L9: 100 mm, L10: 200 mm, L12: 30 mm, (all AC254-XXX-A-ML, Thorlabs), SL: 50 mm (Leica), TL: 200 mm (Leica), OBJ: HC PL APO 100x/1.40 Oil STED White (15506378, Leica). **Filters**: BPF1: ET705/100m (Chroma), BPF2: ET615/30m (Chroma), NF1: NF03-785E-25 (Semrock), NF2 and NF3: ZET785NF (Chroma), CUF: CT780/20bp (Chroma). **Dichroic mirrors**: DM1: T610spxxr (Chroma), DM2: T700dcspxruv_UF3 (Chroma), DM3: ZT405/488/561/640/775rpc (Chroma), DM4: T860SPXRXT (Chroma), DM5: ZT514RDC (Chroma), DM6: Di02-R635 (Semrock). **Lasers**: 561: PDL561 (561 nm, Abberior Instruments), 640: LDH-D-C-640 (640 nm, PicoQuant), 775: KATANA 08 HP (775 nm, OneFive), 980: CP980S (Thorlabs). **Detectors/cameras**: APD1: SPCM-AQRH-13-TR (Excelitas), APD2: SPCM-AQRH-14-FC (PerkinElmer), CMOS: DMK 33UP1300 (The Imaging Source Europe GmbH, Bremen, Germany). **Scanners**: GX/GY: 6215H Galvanometric mirrors + 71215HHJ 671 Servo Driver (Cambridge Technology), PZ: Z-piezo stage LT-Z-100 (Piezoconcept, Lyon, France), XY-stage: SCAN IM 130 x 85 – 2 mm (Märzhäuser Wetzlar GmbH, Wetzlar, Germany). **Fiber optics**: F1: PMJ-3AHPM3S-633-4/125-3-3-1 (OZ Optics), F2: P5-488PM-FC-2 (Thorlabs), F3: M31L01 (Thorlabs), FC1: 60SMS-1-0-A2-02, FC2: 60SMS-1-0-A18-02, FC3: 60SMS-1-4-M5-33 (Schäfter + Kirchhoff GmbH, Hamburg, Germany). **Misc.**: PH: P75H (Thorlabs), AP: D15S (Thorlabs), PBS: PTW 1.15 (Bernard Halle Nachfl. GmbH, Berlin, Germany), λ/4-1: 600-1200 achr. (RAC 5.4.15, B.Halle Nachfl.), λ/4-2: 460-680 achr. (RAC 3.4.15, B.Halle Nachfl.), λ/4-3 and λ/4-4: 500-900 achr. (RAC 4.4.15, B.Halle Nachfl.), λ/2: 500-900 achr. (RAC 4.2.15, B.Halle Nachfl.), SM: 50 mm concave mirror (CM508-050-P01, Thorlabs), SLM: LCOS-SLM X10468-02 (Hamamatsu Photonics), AOM: MT110-B50A1.5-IR-Hk + MDS1C-B65-34-85.135-RS (AA Opto Electronic), M: BB1-E02 (Thorlabs), all other mirrors: PF10-03-P01 (Thorlabs).

### 2.2 Focus lock, tiling and microscope control

An infrared laser (CP980S, Thorlabs), coupled into the objective with a dichroic mirror (T860SPXRXT, Chroma Technology) between the scan and tube lens, is aligned to reflect off the coverslip through total internal reflection. The reflected laser beam is separated from the incoming beam through a D-shaped mirror, and imaged onto a CMOS camera (DMK 33UP1300, The Imaging Source Europe). Through a feedback loop reading the position of the laser beam on the camera chip and subsequently moving the sample with a piezo element in the z-dimension (LT-Z-100, Piezoconcept), the focus lock system keeps the sample in the desired focal plane during shorter and longer acquisitions.

The microscope is controlled through two separate software; image acquisition and most hardware control is done through the Imspector software (Max-Planck Innovation, Göttingen, Germany) while the rest of the hardware control (SLM, focus lock and 775 nm laser output power) is done through Python-based custom-written microscope control software Tempesta (https://github.com/jonatanalvelid/Tempesta-RedSTED, adapted from https://github.com/TestaLab/Tempesta)[7,16]. Line-by-line image acquisition for the two channels is achieved by sequential illumination of each line with two excitation lasers, alternating the detector read-outs and matching the STED power for each fluorophore with the AOM. The 775 nm laser acts as the master trigger, pulsing at 40 MHz, and triggers the two excitation lasers through two separate picosecond delayers (PSD-065-A-MOD, Micro Photon Devices). This allows for picosecond level control of the delay between the excitation pulses and depletion pulses to optimize the STED efficiency and hence resolution and image quality for specific fluorophores.

The pixel size was set to 20–30 nm. Two-colour STED images were recorded as a line-by-line scan for the two channels with a dwell time of 20 µs for each channels.

The tiling is done through custom-written Python-based control in Tempesta of the mechanical stage (SCAN IM 130 × 85 – 2 mm, Märzhäuser). During the tile-scanning, the stage is automatically moved to the next tile after detecting an input signal when the current tile has finished scanning. During the moving of the mechanical stage, the focus lock is automatically unlocked and again locked at the new tile position in the same focal plane. The recorded tile images are stitched together in ImageJ using the Grid/collection stitching plugin[17].

### 2.3 Hippocampal neuron cultures

Primary hippocampal neurons were prepared from embryonic day E18 Sparague Dawley rat embryos. The hippocampi were dissected and mechanically dissociated in MEM. The cells were seeded on poly-D-ornithine (P8638, Sigma-Aldrich, St. Louis, MO, USA) coated #1.5 18mm (Paul Marienfeld GmbH, Lauda-Königshofen, Germany) glass coverslips in 12-well plates at a concentration of 20 × 10^3^ cells per well. They were let to attach to the coverslips with 10% horse serum (26050088, Thermo Fisher Scientific, Waltham, MA, USA), 2mM L-Glut (25030-024, Thermo Fisher Scientific), and 1mM sodium pyruvate (11360-070, Thermo Fisher Scientific). The media was changed after 3h with Neurobasal (21103-049, Thermo Fisher Scientific) supplemented with 2% B-27 (17504-044, Thermo Fisher Scientific), 2mM L-glutamine and 1% penicillin-streptomycin. The neurons were fixed and labelled according to the protocol described under Sample preparation, and experiments were performed on mature neuronal cultures.

### 2.4 Sample preparation

Primary hippocampal neurons used for the imaging were fixed using 4% PFA in PBS for 10 min, permeabilized by incubation with 0.5% TRITON™ X100 in PBS for 5 min, and blocked with 5% BSA in PBS for 30 minutes, all at room temperature. Following this, the cells were incubated with a mixture of the primary antibodies in BSA in PBS for 1 hour, incubated with a mixture of the secondary antibodies in BSA in PBS for 1 hour, and later mounted in Mowiol mounting medium. Washing with PBS was performed between each step.

Gold beads samples were prepared by using a solution of gold beads (80 nm) in water, poly-L-lysine and Mowiol mounting media. First, a layer of poly-L-lysine (P8920, Sigma-Aldrich) was put on a coverslip, incubating it in room temperature for 10 min, then rinsing off the excess solution. Then, the gold bead solution was put on the cover slip, again incubating for 10 min in room temperature and rinsing off the excess solution. Finally, a few µl of Mowiol was put on the cover slip and this was then sealed with the glass slide.

The two-colour labelled DNA origami nanorulers for STED (STED 140ROR, GATTAquant GmbH, Hiltpoltstein, Germany) are nanorulers with three pools of dyes along the length.

### 2.5 Image analysis

The DNA origami nanoruler images were automatically analysed with a MATLAB script. The script loads the two-channel image and detects all the peaks in the 2D images thresholding away the uniform background. Four distinct line profiles at 0°, 45°, 90° and 125° angle to the x-axis respectively, are taken across each detected spot of a ruler. Each line profile is fit with a single Lorentzian function, and the FWHM extracted from the fit coefficient values if the R-square value of the fit R^2^ > 0.9. The final FWHM of the detected spot of a ruler is calculated as the mean value of the two smallest values of the successful fits.

## 3. Results

### 3.1 Tile-scanning STED nanoscopy platform with large FOV

In STED nanoscopy the ability to record homogeneous FOVs in terms of contrast and spatial resolution is tightly connected to the STED and excitation beam co-alignment. Therefore, our first goal was to develop a scanning system generating back-aperture-stationary beams (figure 1).

**Figure 1.**
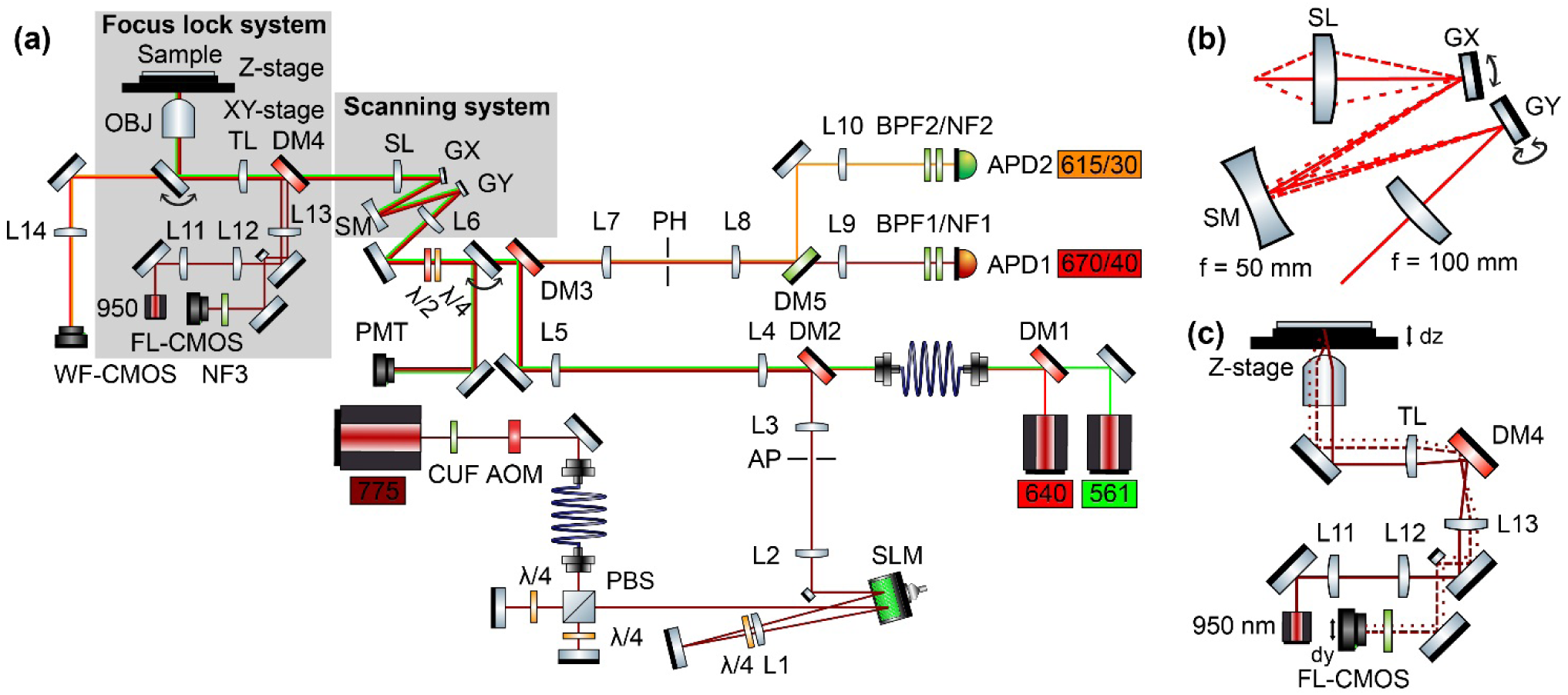
Schematic of the STED setup. (a) Drawing of the setup. Named elements are: L, lenses, DM, dichroic mirrors, BPF, bandpass filters, NF, notch filters, CUF, clean-up filter, APD, avalanche photodiode detectors, PMT, photomultiplier tube detector, SM, spherical mirror, G, galvanometric mirrors, SL, scan lens, TL, tube lens, OBJ, objective lens, λ/2, half-wave plates, λ/4, quarter-wave plates, AP, aperture, PH, pinhole, AOM, acousto-optic modulator, SLM, spatial light modulator, and 561/640/755, picosecond pulsed lasers. A thorough description of the setup and every named element can be found in the Materials and Methods. (b) Zoom-in of the scanning system marked in (a), including an f = 100 mm lens, an X-scanning galvanometric mirror, a spherical mirror, a Y-scanning galvanometric mirror and a scan lens. The solid line shows the aligned beam. The dotted and dashed lines show two extreme scanning positions of the galvanometric mirrors. (c) Zoom-in of the focus lock system marked in (a), including a 950 nm laser, a 150 mm lens, a 30 mm lens, a 200 mm lens, the tube lens, the objective lens, the XY-stage, the Z-stage, the sample, a notch filter, a CMOS camera and dielectric mirrors. The solid line shows the incoming beam, while the dotted and dashed lines show the reflected beam for two different z-positions of the sample.

The scanning system creates the back focal plane stationary beams by utilizing two galvanometric mirrors, with the first imaged onto the second using a spherical mirror (figure 1(b)) and subsequently both imaged onto the back focal plane by the scanning and tube lens. With this system, the whole beams are ensured to enter the objective lens back aperture to avoid vignetting. Instead, the scanning in the focal plane is created by varying the angle of incidence to the back aperture plane. By not cutting the beams in the back aperture, the shape of the PSFs in the focal plane, especially for the doughnut-shaped depletion beam, stay more homogeneous over the whole FOV than when using a scanning system where the beams are both moving and changing the angle of incidence to the back aperture, such as a close-coupled galvanometer mirror system. Moreover, the spherical mirror allows an achromatic focusing of the scanned beams between the galvanometric mirrors, preventing any chromatic aberration in the scanning angles applied to the beams and results in co-aligned beams across a large 80 × 80 um^2^ FOV.

The focus lock is based on a feedback loop system that works on the principle that an infrared beam is coupled into the objective lens as close to the edge of the back aperture as possible, thereby reflecting off the sample by total internal reflection and again captured by the objective lens. This signal is then shifted in the beam path as compared to the incoming beam and hence decoupled by using a D-shaped mirror and then imaged onto a camera. When the sample moves along the optical axis (figure 1(c)), thereby changing the effective focal plane of the excitation and depletion laser beams, the reflection of the infrared beam will move further off the path of the incoming beam with respect to the previous sample position. This will cause a shift of the beam position on the camera sensor. By recording a constant stream off the camera and analysing the frames, the spot can be localized, and a response can be sent to the z-piezo to move the sample in the opposite direction. Characterization of the system yields a sensitivity of 30 nm per pixel on the camera, and a capability of keeping the infrared beam on the camera inside ±0.5 pixels at all times. Thus, the feedback loop will efficiently keep an accurate focal plane during the recording of any STED image with the system. The controlling software widget, integrated in our custom-developed Tempesta software, is also capable of automatically recognizing steps taken in a z-stack recording and any movement that occurs when the mechanical stage moves the sample from one tile to another in a tiled recording, and will re-lock the focus in the focal plane.

The mechanical stage is also controlled through a custom-written tiling widget in the Tempesta software, allowing for manual movement of the stage and an automatic tiling mode where the number of tiles, size of tiles and tile overlap can be arbitrarily decided. Once the tiling has started, the software waits for an input signal when the recording of the current tile has finished in order to move on to the next tile. While moving to the next tile, the focus lock is temporarily unlocked and subsequently locked to the same focal plane.

### 3.2 Characterization of the FOV

The quality of the single-frame FOV was characterized both by imaging gold beads (figure 2) and two-colour DNA origami nanorulers (figure 3). The gold beads images show the co-alignment of the beams (561 nm, 640 nm and 775 nm) at the centre and all four corners of an 80 × 80 µm2 FOV (figure 2(a,b)). Furthermore, a line profile through the beads shows the co-alignment and a background-limited zero of the doughnut (figure 2(c)). The doughnut beam looks homogeneous over the entire FOV, with a central zero intensity measured in the gold beads reflection as below 1% of the crest value across all regions, thanks to the back-aperture-stationary beam scanning system allowing the whole beam to enter the objective lens no matter where in the FOV the beam is scanning. Together, this confirms that this scanning system allows big FOV imaging. Still, minimal chromatic shift is detectable at the corners, but without significantly affecting the measured spatial resolution.

**Figure 2.**
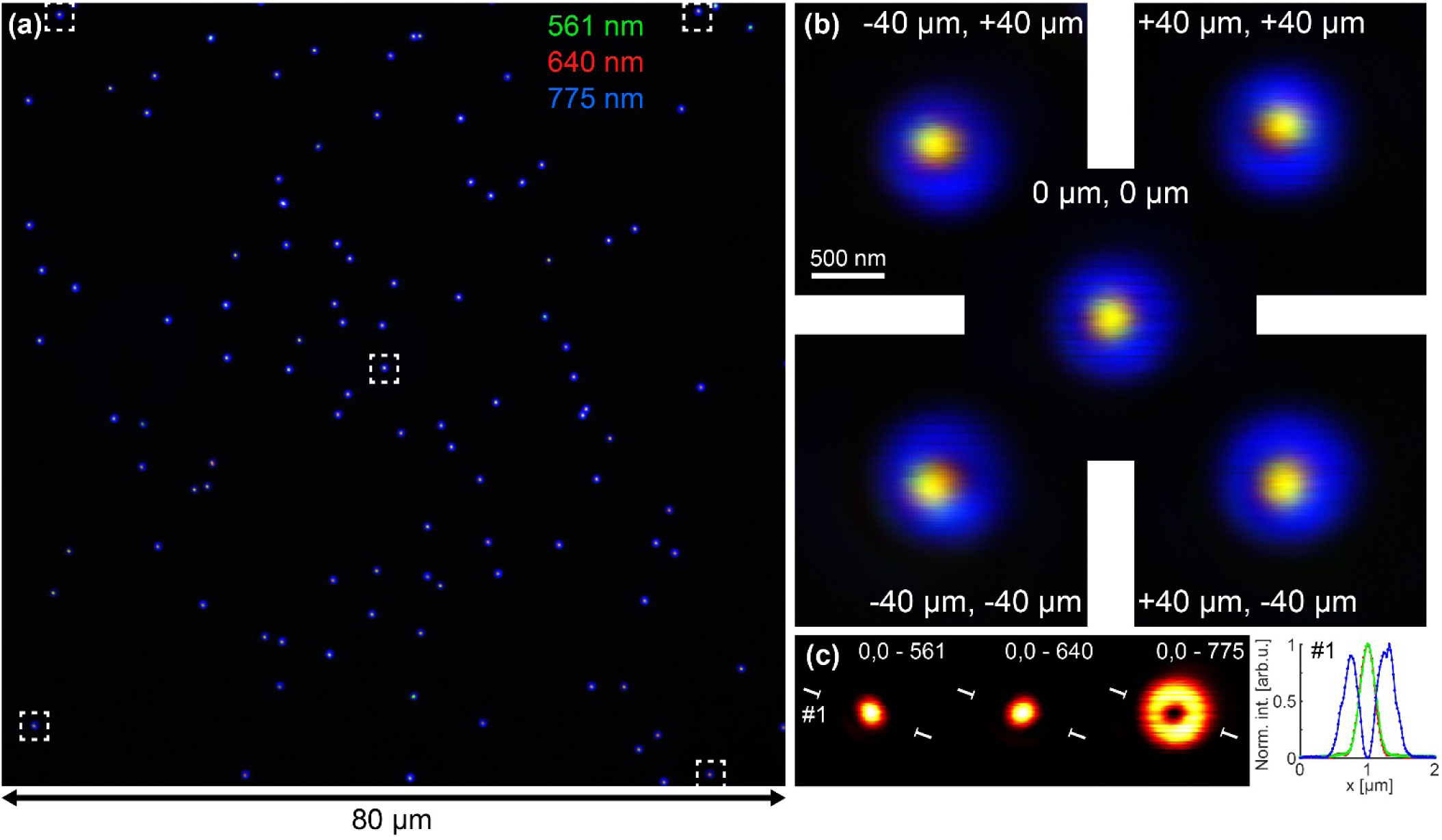
PSF alignment over the field-of-view. (a) 80 × 80 µm^2^ large FOV with the 561 nm (green), 640 nm (red) and 775 nm (blue) laser beams scanned over a sample of gold beads, detected with a PMT. (b) Zooms of single beads in the centre (X0, Y0), top left (X-, Y+), top right (X+, Y+), bottom left (X-, Y-) and bottom right (X+, Y-) corners, as marked in (a). (c) The centre bead (c) in (a) split into the three separate channels of 561 nm excitation beam, 640 nm excitation beam and 775 nm depletion beam. The line profile show the one dimensional intensity profiles of the focal plane of the beads for the two excitation (green, red) and depletion (blue) beams.

**Figure 3.**
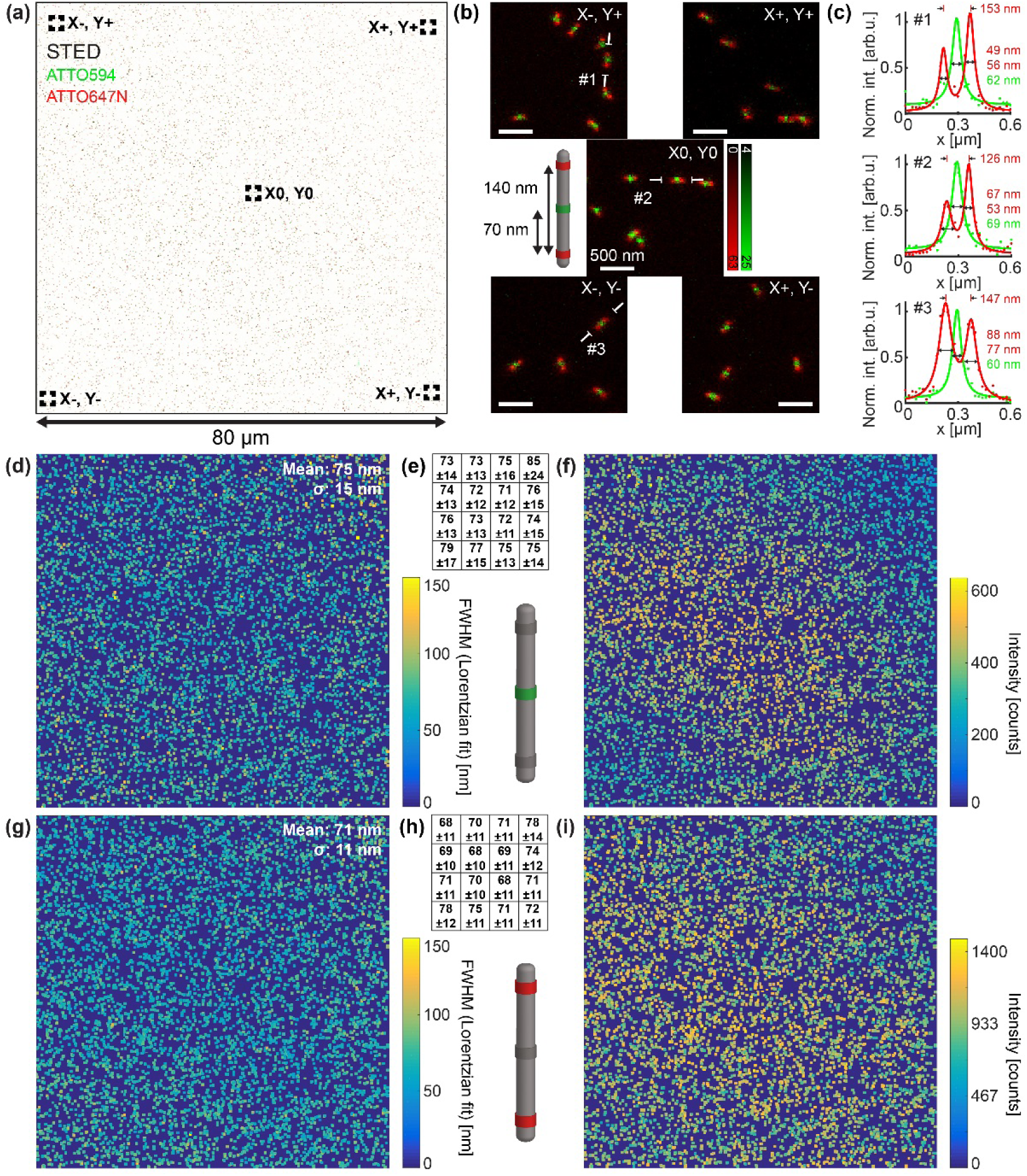
Characterization of the spatial resolution and fluorescence intensity measured over the field-of-view with DNA origami nanorulers. (a) STED image of two-color DNA origami nanorulers distributed in a square area of 80 × 80 µm^2^ large FOV. The pixel size is 20 nm. (b) Zooms of regions with nanorulers in the center (X0, Y0), top left (X-, Y+), top right (X+, Y+), bottom left (X-, Y-) and bottom right (X+, Y-) corners, as marked in (a). (c) Line profiles of a nanoruler from the top left (#1), center (#2), and bottom left (#3) regions of the FOV, as marked in (b). Dots are raw data points, and lines are fitted Lorentzian curves. (d,f,g,i) Colour-coded maps of all detected nanorulers FWHM extracted from a Lorentzians fit (d,g) or integrated intensity (7×7 pixels around the detected center) (f,i), for the ATTO594 (d,f) and ATTO647N (g,i) parts of the nanorulers respectively. (e,h) A schematic map of the FOV shown in (d) and (g) split-up into 16 regions, with the respective mean FWHM and standard deviation in nanometers for each squared 20 × 20 µm^2^ region across the 80 × 80 µm^2^ FOV.

For testing resolution and uniformity of the FOV on a multi-colour fluorescent sample, DNA origami nanorulers make the perfect sample. Thanks to the homogeneity of the nanorulers in signal and size and the flatness of the sample, we can compare and demonstrate the STED imaging quality over the 80 × 80 µm^2^ FOV (figure 3). The whole FOV (figure 3(a)) show nanorulers with ATTO594 in the middle and ATTO647N on both ends, with the centres of the pools of the dyes spaced 70 nm apart. Looking closer at five individual FOVs, again in the centre and at each corner, we can appreciate the uniform resolution of the images attained from each part of the large FOV (figure 3(b)). Taking line profiles from these FOVs and fitting Lorentzian functions to each one of them, we can see that the spatial resolution is comparable and therefore independent of the location of the nanorulers in the FOV (figure 3(c)).

To further investigate the achievable spatial resolution in the whole FOV we measured the size of each nanoruler in the image by individually fitting Lorentzian profiles to every detected bright spot. Mapping the full-width-at-half-maximum (FWHM) values of the fitted curves with a colour-coded look-up table, we can easily appreciate the uniform spatial resolution across the whole FOV (figure 3(d,g)). The mean FWHM for the ATTO594 centre over the whole FOV is 75±15 nm (figure 3(d)). Splitting up the FOV into 16 individual squares, and separately analysing these, we arrive at mean a FWHM ranging between 71–85 nm, with very similar standard deviations (figure 3(e)). Similarly, for the ATTO647N dyes at the edges of the nanorulers, we quantify a mean FWHM of 71±11 nm (figure 3(g)), with the individual sub-squares showing FWHMs ranging between 68–78 nm, again with very similar standard deviations (figure 3(h)). This also shows that the resolution of the two separate spectral channels is comparable. We can also see in both channels that there are plenty of nanorulers that are showing a FWHM down below 50 nm. Moreover, we can compare the recorded intensity of the detected spots (figure 3(f,i)), and see a mapping with a slight decrease in signal in the top-right and bottom-left corners. The fact that we still have a uniform resolution indicate that this does not arise from a deterioration of the depletion beam quality, but instead arises from the slight chromatic shifts observed at these parts of the FOV, where the excitation beams are slightly shifted outwards as compared to the depletion beam.

### 3.3 Extended sample region tile-scanning STED imaging

By performing beam scanning in the 80 × 80 µm^2^ FOV and further controlling the mechanical stage to sequentially record multiple images in a mosaic fashion, we can acquire STED images of theoretically unlimited sample areas (figure 4), all done automatically. This can be important for highly polarized cells such as neurons, with neurites often branching out hundreds of micrometres from the soma. By automatically acquiring a set of tiles and recording an image covering whole neurons we can analyse cell-to-cell variations of individual neurons. We see here two neuronal soma in close proximity at the centre of the 0.53 × 0.53 mm^2^ sample region, with a third neuron in the top left corner (figure 4(a)). Phalloidin is here used to image the actin cytoskeleton, and thanks to the higher resolution of the STED image we can resolve fine actin structures across the whole FOV (figure 4(b)), such as actin bundles, membrane periodic skeleton (MPS) structures, many close-by filaments around the soma, multiple neurites crossing showing the actin organization in the MPS, a neurite with the MPS visible twisting 180 degrees around itself, as well as an intricate network of small actin filaments (figure 4(b)). A line profile across a part of a filament network show that the STED image reveal four filament spaced 80–160 nm apart and with diameters as small as 40 nm as compared to the confocal image, a line profile across the MPS shows the known 180 nm periodicity, and a line profile across three filaments in close proximity again shows filaments as small as 42 nm (figure 4(c)).

**Figure 4.**
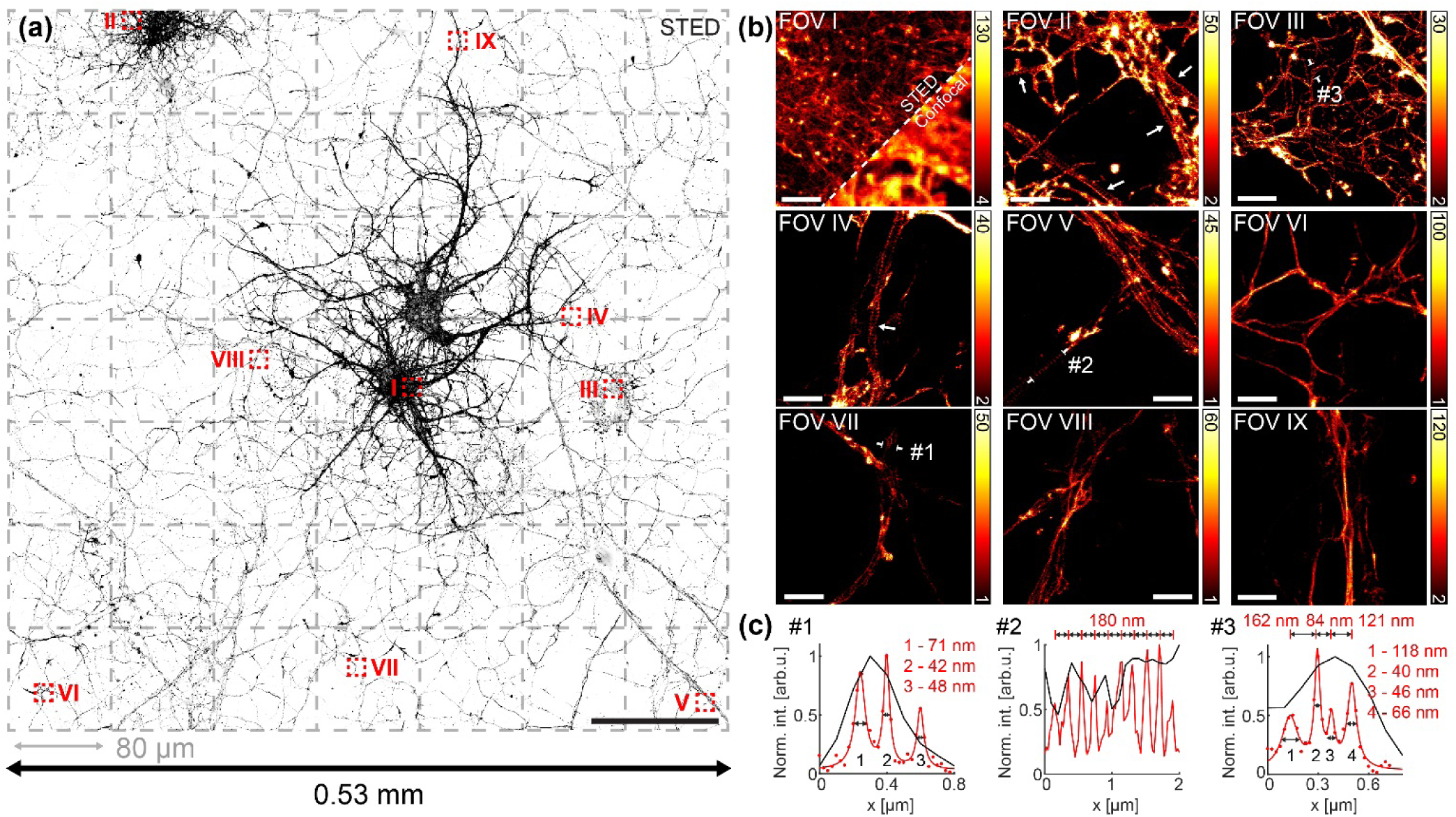
Tile-scanning STED image of hippocampal neuronal culture. (a) 0.53 × 0.53 mm^2^ large FOV of actin stained neurons with Phalloidin_STAR635P, stitched together from 49 individual square tiles, each one 80 × 80 µm^2^ large. The pixel size is 25 nm. Scale bar is 100 µm. (b) Zooms of the regions marked in red boxes in (a), showing STED images of fine actin structures not detectable in the confocal images, such as a group of filaments in the soma (FOV I), the membrane-periodic skeleton (MPS, FOV II), a twisting MPS-striped neurite (FOV III), and many filaments in close proximity (FOV IV), such as three of sizes between 40-66 nm shown in the line profile. Scale bars are 2 µm. (c) Line profiles #1-3 as marked in (b). In #1 and #3, dots in are raw data points, and red lines are fitted Lorentzian curves. In #2, the red line is the raw data. Black lines are confocal data.

## 4. Conclusions

In this work, we introduced a straightforward and fully automated tile-scanning STED implementation to record large, theoretically unlimited-sized FOV without compromising the achievable spatial resolution. We planned and built a microscope with a focus lock, a scanning system and a mechanical stage that together allow imaging of sample regions as big as 0.53 × 0.53 mm^2^, with a single-tile FOV of 80 × 80 µm^2^ with uniform image quality all over.

We show the uniformity at the PSF-scale by scanning a 80 × 80 µm^2^ image of gold beads with all three beams, which demonstrates that the quality of the individual beams are not severely changed at different positions in the FOV. In addition, we recorded a sample of DNA origami nanorulers labelled with Atto594 and Atto647N and showed, in both channels and across the whole FOV, homogeneous FWHMs within 50–80 nm as shown by the automatic fitting analysis.

The imaged sample area was further extended by automating a mechanical stage for tile-scanning STED, approaching the millimetre scale without compromising the spatial resolution. Since the tile recording comes at the expense of time it is important to have a stable and automated microscopy platform, which in our system is achieved with a custom-written Python software controlling both the mechanical stage and a closed-loop focus lock. Future implementations can speed up the recording with smart and specimen-adaptive scanning solutions as in smart RESOLFT[18].

Tile-scanning STED bridges the gap between high resolution studies, which are typically performed in relatively small FOVs showing fine spatial details, and the need to record and analyse big regions of the sample for multi-cellular and tissue-wide studies. For example, neuronal imaging often requires the recording of large portions of the sample since neurons are highly polarized cells with neurites spreading and connecting across larger regions. A whole-nerve-cell study of specific fine structures would then require to have STED spatial resolution across several hundreds of micrometres. Here, we showed the recording of several neuronal cells spread across up to half a millimetre squared FOV in actin-labelled cells.

## Acknowledgements

This work was supported by grant ERC-Stg-638314 from European Commission to I.T.

